# PEPCy: Photostable fluoromodules for live cell, super-resolution microscopy of surface proteins

**DOI:** 10.1101/2024.07.03.601615

**Authors:** Moeka Sasazawa, Afroze Chimthanawala, Rui Zeng, Danah Kim, Katherine Buchan, Ming Zhang, Saumya Saurabh

**Author notes:** Authors’ contributions M.S. and A.C. contributed equally.

## Abstract

We report the evolution and application of two genetically encoded tags that bind the cyanine dyes Cy3 or Cy5 with high specificity and selectivity, in addition to enhancing their photostability. These tags, which we call PEPCy, can be used to target membrane proteins such as G-protein coupled receptors. Due to their orthogonality and high binding-affinity for cognate cyanine dyes, the PEPCy tags can be used for wash-free labeling of cell surface receptors to observe their dynamics at a single molecule level. Together with self-labeling tags, these photostability enhancing proteins against cyanine dyes present a novel, complementary and powerful approach to explore protein dynamics with high spatiotemporal resolution.

## Main

The need for reliable and durable fluorescent markers for live cell microscopy is ever-increasing, driving significant advancements in the development of photostable dyes [1]. Genetic approaches for targeting these dyes are crucial for achieving high-resolution imaging and long-term observations of cellular processes without the detrimental effects of photobleaching [2]. Accordingly, systematic efforts have been made to develop highly photostable rhodamine and cyanine dyes. For example, incorporating azetidine rings into rhodamine dyes enhanced their brightness and photostability, and modifying the azetidine substitutions allowed for fine-tuning of spectral properties and cell permeability [3, 4]. More recently, the lactone to zwitterion transitions in rhodamines has been systematically modulated via auxochrome substitutions further improving their properties for multiple microscopy applications [5]. Modification of cyanine dyes, which have been widely used in microscopy, have improved key parameters such as water solubility, and specificity of cell surface labeling over the decades [6]. More recent work on cyanine dye photostability enhancement has relied on modulating the dye environment, or proximal conjugation of protective agents such as cyclooctatetraene, 4-nitrobenzyl alcohol or Trolox [7–9]. While the mechanism underlying the photostability of these dyes has been debated, the conjugation of protective agents is believed to reduce the population of excited states that lead to photobleaching. [10–12] Despite their superior solubility, cell impermeability, spectral properties, and relatively low cost, applications of cyanine dyes to study protein dynamics have remained limited due to a lack of robust methods to target these dyes to proteins of interest. Currently, haloalkane and benzylguanine conjugated derivatives of cyanine dyes can be covalently targeted via HaloTag (HT) or SNAPTag, although haloalkane and benzylguanine conjugates are neither low cost nor widely accessible [13]. We sought to fill this gap and improve the photophysical properties of off-the-shelf sulfonated cyanine dyes by developing specific tags relying on non-covalent binding to single chain variable fragments (scFv) of human antibodies. scFvs have been utilized to develop fluorogen activating proteins that bind non-covalently to malachite green and thiazole orange and tune their spectral properties and enhance photostability [14]. New fluorogens have been added to the repertoire of fluorogen activating proteins [15–17] and fluorogen activating proteins have been applied for live-cell super-resolution microscopy [18]. Unlike the irreversible binding between organic dyes and covalent tagging schemes, scFvs exhibit tunable binding-affinities for their cognate small molecule, thereby providing photostability enhancement via encapsulation or exchange [19].

Here we performed directed evolution on a library of scFvs to isolate two unique proteins that can bind sulfonated derivatives of Cy3 or Cy5, with high affinity and selectivity. We refer to these tags as photostability enhancing proteins against cyanine dyes (PEPCy) as they enhance the lifetime, molecular brightness, and photostability of their cognate cyanine dyes significantly compared to the same dyes targeted via the self-labeling enzyme, HT. We demonstrate the utility of PEPCy tags in single molecule microscopy and live cell imaging of surface receptors in bacterial and mammalian cells. Together, the orthogonal PEPCy3 and PEPCy5 tags open a new paradigm that enables simultaneous multi-color single molecule localization microscopy at superior spatiotemporal resolution for imaging surface proteins using cyanine dyes.

Given the accessibility of cell impermeant, sulfo-cyanine dyes and the success of scFv yeast surface display libraries in generating fluorogen activating peptides, we screened for binders of these dyes. Directed evolution was performed on a yeast cell surface display library of diverse scFvs (Fig. 1A and S1A). In addition to modulating the concentrations of Cy3 or Cy5 to obtain high affinity scFvs, we also applied a photobleaching selection pressure to select for the most photostable variants. A stringent selection was applied at each step, collecting yeast cells exhibiting top 0.1% of the population in their respective fluorescent channels (Fig. S1B, C) Characterization of spectral properties of cognate dyes bound to purified PEPCy tags revealed approximately a six-fold enhancement in molecular brightness and bathochromic shifts of 12 nm (excitation) and 6 nm (emission) for the PEPCy3-Cy3 complex relative to Cy3 alone (Fig. 1B and S1D, Table 1). The PEPCy5-Cy5 complex exhibited a meagre 7% enhancement in molecular brightness and no spectral shifts (Fig. 1C and S1D, Table 1). Critically, each PEPCy tag exhibited high specificity for its cognate cyanine dye, and incubation of cells expressing PEPCy3 with Cy5 or vice versa did not lead to cross reactivity in microscopy experiments (Fig. 1D). On the yeast cell surface, PEPCy3 bound Cy3 with a *K_D_* of 8 nM while PEPCy5 bound Cy5 with a *K_D_* of 80 pM (Fig. S1E, F, Table 1). Due to the innate environmental sensitivity of cyanine dyes [20, 21], we explored whether PEPCy tags modified the fluorescence lifetimes of their cognate dyes. Fluorescence lifetime microscopy (FLIM) of PEPCy expressing yeast cells showed that both PEPCy3 and PEPCy5 increased the lifetime of their cognate cyanine dyes compared to covalent targeting of the dyes via HT (Fig. 1E and S1G, H). Well separated lifetimes can be utilized for differentiating between a mixed population of cells expressing PEPCy or HT in the same spectral window. Having characterized the biochemical and spectral properties of PEPCy fluoromodules, we next evaluated their utility in single molecule localization and time lapse microscopy of cell surface proteins.

**Fig. 1:**
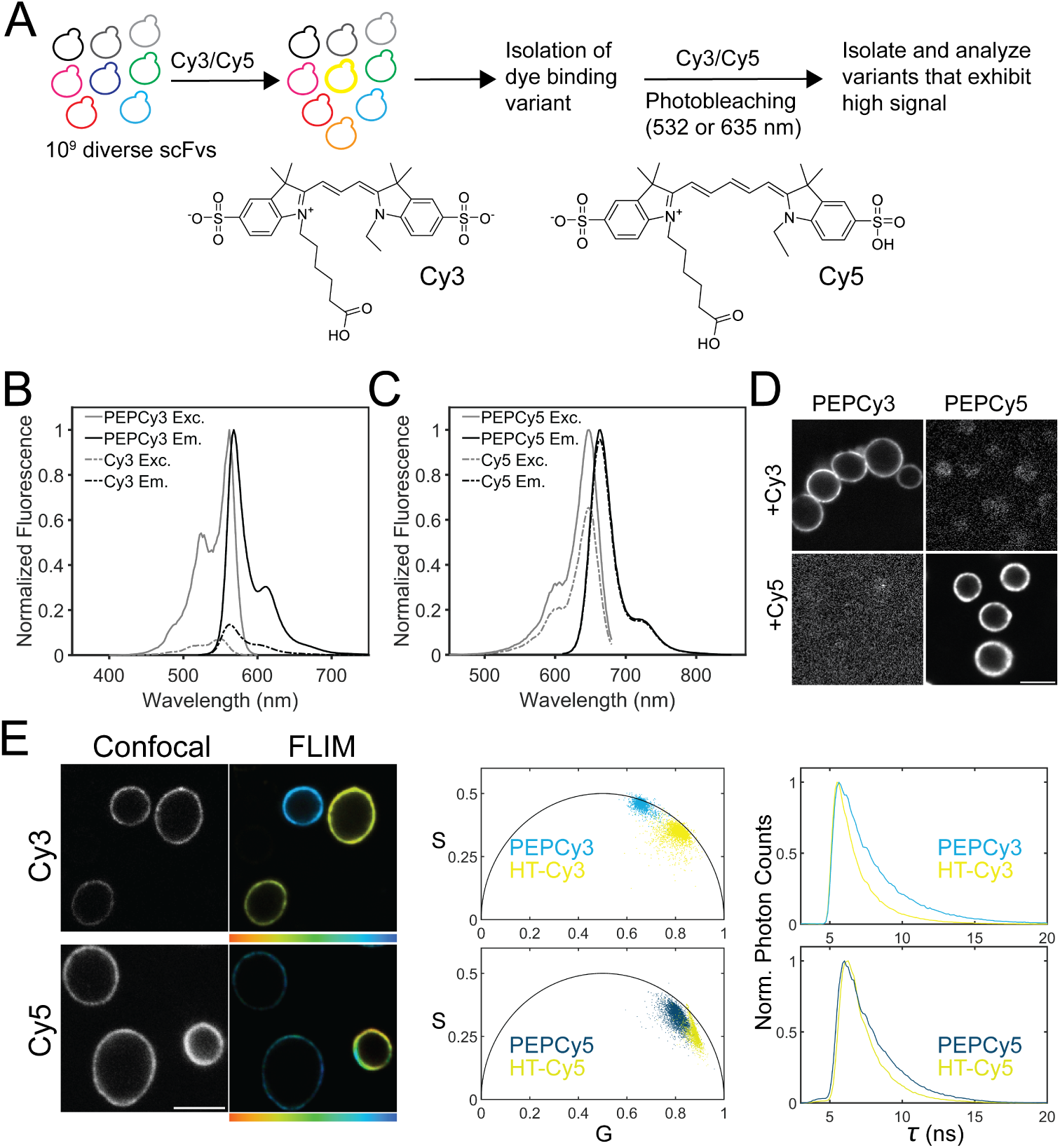
Selection and spectroscopic characterization of PEPCy tags. **A**. Schematic of the directed evolution of scFvs with a yeast cell surface display. Structures of the cyanine dyes, Cy3 and Cy5. Excitation and emission spectra for, **B** free Cy3 and PEPCy3 complex, and **C** free Cy5 and PEPCy5 complex. For complexation, 100 nM dyes were incubated with 1 *µ*M PEPCy proteins for 30 mins before obtaining spectra. **D**. Confocal images showing yeast expressing (left) PEPCy3 and (right) PEPCy5, incubated with either Cy3 (top) or Cy5 (bottom). Images of cells with non-cognate dyes are scaled so as to discern the background clearly. **E**. (left) Confocal images of a mixed population of labeled cells expressing (top) PEPCy3(Cy3) and HaloTag(Cy3-HTL), and (bottom) PEPCy5(Cy5) and HaloTag(Cy5-HTL), respectively. (right) Fluorescence lifetime images of the same cells. The lifetime colormaps extend from 1 ns to 3 ns (Cy3) and 1 ns to 2 ns (Cy5). Phasor plots and decay curves corresponding to the FLIM micrographs show populations with distinct lifetimes for PEPCy tags compared to HaloTag targeted Cy dyes. All scale bars in the figure are 5 *µ*m.

**Table 1.** Spectral and binding properties of PEPCy fluoromodules and their cognate cyanine dyes.

| Fluor | MW<br>(kDa) | $\lambda_{max}Ex$<br>(nm)<sup>c</sup> | $\lambda_{max}Em$<br>(nm)<sup>c</sup> | EC<sup>a</sup><br>(M<sup>-1</sup>cm<sup>-1</sup>)<sup>c</sup> | QY<sup>b</sup> | MB<br>(M<sup>-1</sup>cm<sup>-1</sup>)<sup>d</sup> | $K_D$<br>(nM)<sup>e</sup> |
| --- | --- | --- | --- | --- | --- | --- | --- |
| Cy3 | 0.63 | 548 | 562 | 150000 | 0.04 | 6 | – |
| PEPCy3 | 31.18 | 558 | 568 | 126923 | 0.27 | 34.3 | 8 |
| Cy5 | 0.66 | 650 | 664 | 250000 | 0.27 | 67.5 | – |
| PEPCy5 | 24.28 | 650 | 664 | 251168 | 0.29 | 72.8 | 0.08 |

<sup>a</sup>Extinction coefficient
<sup>b</sup>Quantum yield
<sup>c</sup>Measured in PBS, pH 7.4.
<sup>d</sup>Product of EC and QY divided by 1000.
<sup>e</sup>Cell surface $K_D$ .

PEPCy tags were expressed on the surface of mammalian cells and imaged by Total Internal Reflection Fluorescence (TIRF) microscopy (Fig. 2A). Human embryonic kidney (HEK293) cell lines transiently expressing the *β*2 adrenergic receptor (B2AR) N-terminally fused with PEPCy3 or cells expressing a point mutant of B2AR (B2AR-Ala) N-terminally fused with PEPCy5 were generated. HT-B2AR expressing HEK293 cells were used for comparison. Upon labeling with their cognate dyes, all samples exhibited single molecule signals on the cell membrane (Fig. S2A, B, supplementary videos 1 and 2). A key difference in the background fluorescence was observed between the non-covalent PEPCy tags and the covalent HT. Both PEPCy3 and PEPCy5 single molecule localizations displayed a higher signal above the background compared to the HT targeted cyanine dyes (Fig. S2A, B). We also observed significantly higher background counts for HT-Cy3 compared to PEPCy3 localizations (Fig. S2C). Due to the reduced cellular autofluorescence in the Cy5 spectral window, we observed a small but significant decrease in the background counts from PEPCy5 compared to HT-Cy5 (Fig. S2D). Notably, PEPCy fusion proteins could be labeled using 10 pM-10 nM dye concentrations within ten minutes while HT labeling required a longer incubation with over 100 nM dye and multiple washes for single-molecule imaging. These improvements in signal to background fluorescence from PEPCy tags can be attributed to their non-covalent, high affinity interactions with their cognate dyes. Owing to its low *K_D_* for Cy5, we could label HEK cells expressing PEPCy5-B2AR-Ala using 10 pM dye followed by TIRF microscopy without washing any unbound dye (Fig. S2E, supplemetary video 3). We observed a decrease in the number of single molecule localizations over 1.5 seconds of imaging followed by turning off the illumination beam for 2 seconds. During this time, new PEPCy5-B2AR-Ala molecules diffused in the field of view. This process could be performed iteratively to track multiple single molecules on the same cell, thereby obviating the need for constantly replenishing or washing the dye.

**Fig. 2:**
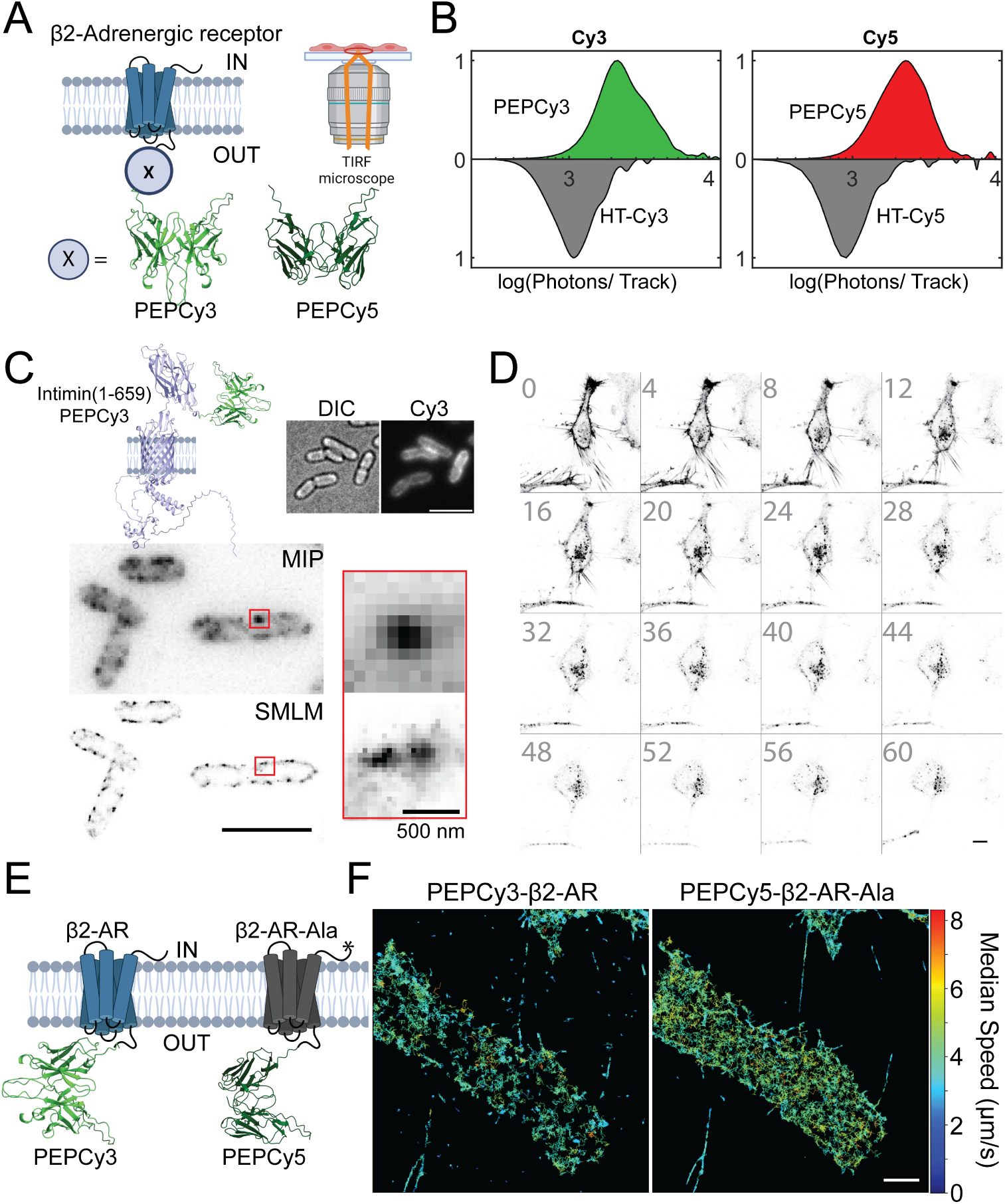
Applications of PEPCy tags for imaging cell surface receptors. **A**. Schematic of membrane protein tracking experiment using TIRF microscopy. AlphaFold predicted structures of PEPCy tags. **B**. Normalized probability distributions of photons detected per trajectory from single particle analyses between PEPCy and HT targeted to their cognate dyes. (N = 1000-1300 single tracks per sample). **C.** Super-resolution microscopy of Intimin fused to PEPCy3 expressed on *E.coli* outer membrane. Schematic of the fusion is shown followed by diffraction-limited DIC and fluorscence images of the cells expressing the same. Maximum intensity projections (MIP) compared with a 2D reconstruction of single molecule localization data. Inset compares a cluster observable in the MIP that is resolved into two clusters in the super-resolved image. **D**. Time-lapse confocal microscopy of HeLa cells expressing PEPCy3-B2AR labeled using Cy3. Numbers on the left of each image denote the time in minutes. Isoproterenol was added at t = 0 and imaged were acquired every 30s for 1 hour. **E**. Schematic of simultaneous two color tracking of PEPCy3-B2AR and PEPCy5-B2AR-Ala on the same cell. **F**. Single molecule trajectories of PEPCy3 (left) and PEPCy5 (right) on a HEK cell, color coded by median trajectory speed. All scale bars in the figure unless mentioned otherwise are 5 *µ*m. A and E were created with BioRender.com.

The number of photons detected by a fluorescent single molecule before it bleaches irreversibly is a robust measure of its photostability. Accordingly, we compared the photons detected per trajectory obtained from adrenergic receptor fusions of PEPCy tags on HEK cell surface with HT targeted Cy3 or Cy5. PEPCy tags exhibited at least a two-fold improvement in the total photon output compared to the cyanine dyes covalently conjugated to HT (Fig. 2B). To exploit the high photon output of PEPCy tags in single molecule localization microscopy (SMLM), we fused the bacterial surface protein Intimin(1-659 residues) to PEPCy3 and labeled the samples with Cy3 (Fig. 2C). Surface associated signal was clearly visible with diffraction-limited microscopy. Maximum intensity projections (MIP) of Intimin-PEPCy3 revealed clusters on the cell surface, which could be further resolved into sub-diffraction structures via single molecule localization, showing the utility of PEPCy3 in SMLM in bacterial cells (Fig. 2C). To demonstrate the utility of the enhanced photostability and molecular brightness of Cy3 or Cy5 bound to cognate PEPCy tags, we performed time-lapse confocal microscopy on HeLa cells expressing B2AR fusions of PEPCy tags. PEPCy3-B2AR cells exhibited a membrane localized signal which upon addition of the agonist, isoproterenol, got internalized over an hour (Fig. 2D, supplementary video 4). Unstimulated HeLa cells expressing PEPCy5-B2AR-Ala remained localized to the cell membrane over 30 mins (Fig. S2F, supplementary video 5).

Traditionally, the comparison of a mutant protein with its WT counterpart relies on expressing the proteins in two different cell lines. However, due to their orthogonality, PEPCy constructs can be used to directly compare WT and a mutant of a cell surface protein. To demonstrate this, we co-expressed a fusion of B2AR with PEPCy3 and a fusion of a B2AR-Ala with PEPCy5 (Fig. 2E). The B2AR-Ala mutant contains a C-terminus Alanine that disrupts the interaction between the receptor and down-stream signaling proteins through a PDZ domain [22]. HEK293 cells co-expressing the PEPCy tagged copies of wildtype and mutant B2ARs were imaged using a TIRF microscope with two detectors capable of spectrally resolving PEPCy3 signal from that of PEPCy5, thereby allowing for simultaneous measurement of diffusing receptors on the cell surface (Fig. 2F, supplementary video 6). Images from the two channels were registered using fiduciaries with a median target registration error of 19.8 nm. Upon analyzing the trajectories of B2AR and B2AR-Ala from the same cells, we found that the mutant protein exhibited a fast moving population on the cell surface compared to the WT protein (Fig. 2F, Fig. S2G). This observation was further supported by mean squared displacement analyses (Fig. S2H). In addition to B2AR, we have demonstrated the applicability of the PEPCy3 tag in super-resolution microscopy of the sodium channel Nav1.7 on trigeminal ganglia neuron cell surfaces, elsewhere [23]. To expand the live-cell labelling toolkit for targeting dyes to proteins of interest, we developed genetically encoded, high-affinity and orthogonal protein tags for two widely used sulfo-cyanine dyes, Cy3 and Cy5 that differ by a single double bond. We have vastly improved the fluorescence properties of Cy3 bound to PEPCy3, bringing this dye close in performance to Cy5, while also bringing Cy3’s excitation maximum closer to a commonly used laser line. Both PEPCy tags improve the molecular brightness and fluorescence lifetimes of their cognate cyanine dyes, that could be used to further distinguish dyes with overlapping or similar spectra. The high affinity of PEPCy tags allows for fast and wash-free labeling with high signal to background compared to covalent tagging schemes. The enhanced photostability of PEPCy tags can be used to study temporal changes over longer time scales in membrane protein trafficking at both an ensemble and single molecules levels. Finally, orthogonal PEPCy tags can be applied for simultaneous single-molecule imaging of at least two cell surface proteins, while being complementary and compatible with other labelling tools that could further increase the number of simultaneously imaged structures.

Despite these advances, PEPCy tags are limited to optimally label cell surface proteins. Future work on modifications of the PEPCy residues will aim at expanding them for cytosolic and organelle specific expression. Subsequent PEP libraries will be developed to bind other dye families such as rhodamines and bodipy. Another untapped opportunity lies in the area of dye modifications. Here, we will focus on optimizing PEPCy tags to target environmentally sensitive[24] and highly photostable cyanine[8] derivatives. A final limitation to be overcome is the elucidation of structural and molecular mechanisms underlying the photostability enhancement of these versatile tags. This will allow for rational design of PEP tags to obtain the desired spectral and photochemical properties for their cognate dyes.

## Methods

### Directed evolution of PEPCy tags

A yeast cell surface display library of scFvs with a diversity of 10^9^ clones was utilized to screen for binders of Cy3 and Cy5 (Fig. S1A)[25]. The library comprised of the plasmid pPNL6 in the *S. cerevisiae* strain JAR200, provided by Andrew Rakestraw (Massachusetts Institute of Technology, Cambridge, Massachusetts, USA), and obtained from Christopher Szent-Gyorgyi (Carnegie Mellon University). Sulfonated Cy3, Cy5 and their chlorohexane conjugates were provided by Dr. Brigitte Schmidt (Carnegie Mellon University). To perform the selection, the yeast cell library elements were induced in 1 L baffled flasks (500 mL growth media, 20 g dextrose, 5 g casamino acids, 1.7 g Yeast nitrogen base, 5.3 g ammonium sulfate, 7.4 g sodium citrate, 2.2 g citric acid) for three days at 20°C prior to being screened for Cy3 or Cy5 binding on a FACSVantage SE flow cytometer with FACSDiva Option (BD biosciences). For selection of scFvs against Cy3, a 532 nm laser was used for excitation while collecting fluorescence through a 575/26 nm band-pass filter and a 530/30 nm bandpass filter to detect Alexa488 conjugated anti-myc antibody for validation in round IV (Fig. S1B). For Cy5 selection, a 633 nm laser excitation laser was used and fluorescence was collected through two bandpass filters 685/35 nm and 780/60 nm (Fig. S1C). This was done to collect as much Cy5 signal as possible, on the flow cytometer. Cells were labeled at saturating concentrations of Cy3 and Cy5 (2 *µ*M) for the first round and at 50 nM dye concentration for the next two rounds of sorting using flow cytometry. Cells exhibiting top 0.1% brightness in the respective dye channels were collected. For the fourth round of sorting, cells were labeled with 50 nM cognate dye followed by a 5 minute photobleaching step using an LED light source (540 nm, or 640 nm, Mouser electronics). The rationale for this selection pressure was that high affinity cyanine dye binders with enhanced photostability would be enriched in the resulting population. Single cells from round 4 were picked, grown and induced in the same manner as above and dye binding was validated using a Tecan Safire 2 plate reader. From the positive strains, the scFvs were isolated, the *V_H_* and *V_L_* fragments were cloned out and expressed separately followed by testing for dye binding via fluorescence. We found that the Cy3 binding scFv composed of two *V_H_* domains, while the Cy5 binding scFv was composed of *V_L_* chains. Hence we introduced Gly-Ser linkers and constructed PEPCy3 and PEPCy5 scFvs. Finally, DNA sequences containing PEPCy3 or PEPCy5 were integrated in a pYD1 vector (Gal1 promoter, Trp selection) and transformed into JAR200 cells. For control experiments and to compare PEPCy tags against a self-labeling tag, HT was cloned into a pYD1 vector followed by transformation into yeast cells.

#### PEPCy3 amino acid sequence

EVQLVESGGDLVQPGRSLRLSCTASGFPFGDYAITWFRQAPGKGLEWVGFIRSKPFGGTTEYAASVRGRFTISRDDSKSIAYLQMNSLKAGDTAVYYCTRFSPFHNDRGVYSRDDAFDIWGQGTMVTVSSGILGSGGSGSAGSGGSGSAGSGGSGSAPGGGEVQLVESGGDLVQPGRSLRLSCTASGFPFGDYAITWFRQAPGKGLEWVGFIRSKPFGGTTEYAASVRGRFTISRDDSKSIAYLQMNSLKAGDTAVYYCTRFSPFHNDRGVYSRDDAFDIWGQGTMVTVSSGIL

#### PEPCy5 amino acid sequence

EIVLTQSPATLSLSPGDRATLSCRTSQSVSHHLAWYQQKPGQAPRLLIYGASNRATGIPDRFSGSGSGTDFTLTISRLEPEDFAVYYCQQSPAFGQGTKVEIKSGILGGGGSGGGGSGGGGSGGGGSEIVLTQSPATLSLSPGDRATLSCRTSQSVSHHLAWYQQKPGQAPRLLIYGASNRATGIPDRFSGSGSGTDFTLTISRLEPEDFAVYYCQQSPAFGQGTKVEIKSGIL

### Plasmid, protein, and strain engineering Bacterial strain engineering and protein expression

For expression of recombinant proteins, PEPCy DNA sequence was cloned in a modified pet21a vector (a gift from Dr. Josef Franke, Carnegie Mellon University) that encodes an N-terminal decahistidine tag, followed by a GST and an HRV3C protease cleavage site before the construct of interest. The resulting plasmids were transformed into the *E. coli* strain Rosetta-gami 2 (DE3) (Novagen) and recombinant proteins were isolated using a previously published protocol for scFv purification[16]. Briefly, following induction with 1 mM IPTG at 20°C, cells were washed in 25 mM Tris (pH 8.0) containing 150 mM NaCl. The washed cells were then pelleted and frozen at –80°C. After thawing in wash Buffer (50 mM Tris, pH 7.4, 500 mM NaCl, 30 mM imidazole, pH 8.0, and 0.015% Triton X-100), cells were lysed by sonication in the presence of DNase, lysozyme, and PMSF. The lysate was clarified by centrifugation at 22,789 *×*g for 45 minutes at 4°C, and the supernatant was incubated with Ni-NTA agarose (QIAGEN) for 2 hours at 4°C. After incubation, the beads were transferred to a column and washed. His6-tagged HRV3C protease was then added, and the mixture was incubated overnight. Fresh Ni-NTA agarose was subsequently added, and the released protein was collected from the column after 1 hour of incubation. For making the *E. coli* constructs for imaging Intimin, the C-terminal region of Intimin (1-659 aa) constituting the SP, LysM and *β*-domains and the secreted D0 Ig-like domain was fused to PEPCy3 and inserted into a pET29b plasmid backbone. This plasmid was transformed into BL21(DE3) cells that were used for single molecule localization microscopy, *vide infra*.

### Cloning of lentiviral vector pLenCMVPM-PEPCy3m-B2AR

DNA fragments encoding a codon-modified PEPCy3 monomer, a flexible amino acid linker “GGSGSAGSGGSGSAGSGGSGSAPGGG”, and the original PEPCy3 monomer isolated from library sorting, were assembled in the order indicated by overlap-extension PCR. Terminal oligos used in assembly added a 5’-XbaI and 3’-AscI site to the product. The recipient plasmid backbone was produced using lentiviral transfer vector “pLenCMVPM-Mars2h-B2AR” (unpublished, confers puromycin resistance) by treatment with restriction enzymes XbaI and AscI. Ligation of like-digested insert resulted in placement of PEPCy3 in-frame between DNA encoding the human Ig-kappa signal peptide, and a fusion linker immediately preceding human Beta-2 adrenergic receptor. Plasmid sequences were verified by Sanger sequencing.

### Cloning of lentiviral transfer vector pLenPGKPM-PEPCy5-B2AR-Ala

Lentiviral transfer vector “pLenti-PGK-KRAS(G12V)”, used for creation of the PEPCy5-B2AR-Ala-hosting lentivirus, was a generous gift from the lab of Dr. Michael Lotze (Hillman Cancer Center, Pittsburgh PA). Intermediate plasmid “pLenti-PGK-Mars1Cy-KRAS” was constructed by inserting a Mars1Cy-KRAS transgene, flanked by 5’-AgeI and 3’-BsrGI restriction sites, into the like-digested viral transfer vector immediately downstream of the PGK promoter. A cloning insert encoding “PEPCy5-B2AR-Ala” was constructed by amplifying, via PCR in separate reactions, DNA encoding the human Ig-kappa signal peptide, a synthetic gene encoding PEPCy5 from E. coli expression plasmid pET21a-PEPCy5, and the open reading frame encoding human Beta-2 adrenergic receptor from plasmid pLenCMVPM-PEPCy3m-B2AR. These fragments were assembled, in the order indicated, by overlap-extension PCR using terminal primers that added an SbfI site to the 5’ end of the amplicon, and appended sequence encoding an extra alanine on the C-terminus of B2AR, followed by a stop codon and BsrGI cloning site. After treatment with restriction enzymes SbfI and BsrGI, then insert was ligated into like-digested plasmid backbone produced from pLenti-PGK-Mars1Cy-KRAS(G12V), which hosts a hygromycin resistance marker. Plasmid was recovered from E. coli transformants generated by transformation of the ligation mixture into E. coli and verified by Sanger sequencing.

### Lentivirus preparation, transduction of HeLa cells and HEK cell transfection

Lentivirus was produced using the PhoenixGP cell line (Nolan Lab, Stanford University) by co-transfection of cells in T25 flask format using lentiviral transfer vector (ONE of pLenCMVPM-PEPCy3m-B2AR or pLenPGKPM-PEPCy5-B2AR-Ala, 10 *µ*g), plasmid pCMVd8.74 (5.0 *µ*g), and pseudotyping plasmid pMG2.g (5.0 *µ*g). Transfection mixtures were prepared in serum-free DMEM containing 30 *µ*L of TransIT-LT1 reagent (Mirus Bio). Media containing transfection mixture was replaced after six hours, and virus particles were allowed to accumulate for 48 hours post-transfection. Infectious media was clarified by passage through 0.45 micron filters and immediately applied to HeLa cells seeded in 60 mm dishes to create single (PEPCy3-B2AR OR PEPCy5-B2AR-Ala) or doubly (PEPCy3-B2AR AND PEPCy5-B2AR-Ala)-expressing cells. Transduced cells were selected in culture media (DMEM with 10% FBS) supplemented with puromycin, hygromycin, or both. Clonal cells were isolated after sorting for Cy3 and/or Cy5-binding properties. HEK293 cells were transiently transfected using TransIT-LT1 reagent (Mirus Bio) as per the manufacturer’s protocol. Briefly, confluent cells in T25 flasks were washed with serum-free DMEM and incubated with 30 *µ*L of the transfection reagent pre-mixed with 10 *µ*g of the respective plasmids. Transfection media was replaced after 6 hours and the cells were allowed to grow for 24 hours while being maintained in puromycin (for PEPCy3) and hygromycin (for PEPCy5) supplemented DMEM and 10% serum. HEK293 cells harboring a HaloTag-B2AR plasmid were constructed by transient transfection as above. The HaloTag-beta2-adrenergic receptor was a gift from Catherine Berlot (Addgene plasmid # 66994).

### Spectral properties and binding measurements

For measuring extinction coefficients, a ten-fold excess of purified PEPCy3 or PEPCy5 were incubated with serial dilutions of Cy3 or Cy5, respectively in PBS at pH 7.4 at 4°C for 1 hour. Relative quantum yield measurements were conducted using Cy3 or Cy5 as reference in PBS at pH 7.4 by the gradient method[6]. For each replicate, five samples were constituted using various dye concentrations in the presence of a ten-fold excess purified protein. The solutions were allowed to complex at 4°C for at least 1 hour prior to reequilibration and measurement at room temperature. Fluorescence spectra were obtained using a SAFIRE II fluorescence plate reader (Tecan) equipped with dual excitation and emission monochromators with a 5.0 nm spectral bandwidth. Emission spectra were integrated using Spectragryph, version v1.2.16.1 (https://www.effemm2.de/spectragryph/index.html), and relative quantum yield was determined via linear fit of the area versus absorbance plots. Dye binding isotherms were generated using JAR200 yeast cells expressing the PEPCy scFvs on their surface as per a privously reported protocol.[26] Yeast cells at an optical density (O.D. at 600 nm) of 0.3 expressing PEPCy tags on their surface were incubated with varying concentrations of cognate dyes at room temperature for 30 mins followed by two washes in PBS. As a control for non-specific binding, JAR200 cells that were not induced to express PEPCy were used. Binding of the dye was verified by fluorescence microscopy and the samples were loaded in a multi-well plate followed by readout of fluorescence at the emission maximum using a SAFIRE II fluorescence plate reader. Background fluorescence was subtracted from the data and the resulting points were fit using a single site, non-cooperative binding model according to the following equation:

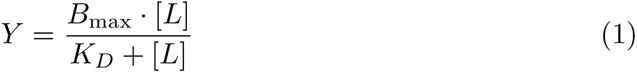

Where, *Y* is the specific fluorescence signal, *B_max_* is the maximum fluorescence signal at saturation, *K_D_* is the binding affinity, and [*L*] is the concentration of dye.

### Cell labeling conditions

#### Yeast labeling

JAR200 strains harboring PEPCy3, PEPCy5 or HT fused to AGA2 were grown overnight in liquid induction media SG+CAA (5.0 g/L casamino acids-TRP, 20 g/L galactose, YNB 6.7 g/L (Difco), 5 g/L (NH4)2SO4, 0.1 M sodium phosphate buffer) at 30°C at 300 rpm. The saturated O/N culture was diluted to an O.D. of 0.1 in induction media and incubated at 30°C, with shaking at 300 rpm. Upon reaching O.D. 0.3, PEPCy3 and PEPCy5 expressing cells were incubated with 100 nM Cy3 or Cy5 dyes, respectively. Labeling was carried out by incubating cells with the dye for 30 mins at 30°C, 400 rpm in a thermomixer. HT expressing cells were labeled either with 100 nM Cy3-Halotag ligand (HTL) or 100 nM Cy5-HTL for 1 hour at 30°C, 400 rpm. After incubation, cells were subjected to centrifugation and the media was discarded. The cell pellet was resuspended in sterile PBS and utilized for imaging.

#### *E. coli* labeling

BL21 cells harboring the Intimin-PEPCy3 were grown in LB at 37°C, 200 rpm to OD 0.1 and incubated with 0.5 mM IPTG for 1 h to induce the expression of Intimin-PEPCy3. Then, a 200 *µ*L aliquot of cells was incubated with 100 nM of Cy3 for 30 min for labelling. Followed by a wash and resuspension in minimal media containing 6 mM Na_2_HPO_4_, 4 mM KH_2_PO_4_, 9.5 mM NH_4_Cl, 0.5 mM MgSO_4_, 0.5 mM CaCl_2_, 0.1% Ferrous chelate solution (F0518, SIGMA), 0.2% glucose). The cells were spotted on a 1.5% agarose pad for imaging.

#### Mammalian cell labeling

For the B2AR trafficking experiments, HeLa cells stably expressing PEPCy3-B2AR on the plasma membrane were plated onto glass bottom dishes 24 hours prior to imaging. Cells were incubated under normal growth conditions (37°C, 5% CO_2_) for 60 minutes prior to imaging. Media was replaced using colorless DMEM (Gibco) supplemented with 100 nM Cy3 added to the culture in the last 15-20 mins followed by replacing the media with colorless DMEM. HeLa cells stably expressing PEPCy3-B2AR were exposed to 20 *µ*M isoproterenol (iso) for 2 mins to initiate receptor internalization. Iso was washed out followed by collection of time-lapse data every 30 seconds for 1 hour. Cells expressing PEPCy5-B2AR-Ala were labeled using 100 nM Cy5 for 20 mins followed by replacing the media. For single molecule tracking of PEPCy3-B2AR or PEPCy5-B2AR-Ala on transiently expressing HEK293 cells, 1 nM or 10 pM of Cy3 or Cy5 was added to the imaging media (colorless DMEM) followed by microscopy.

### Microscopy

#### Confocal and lifetime microscopy of yeast cells

Confocal microscopy was performed on Nikon Eclipse Ti2 microscope using a 100x (1.45 N.A.) oil immersion objective and a quad beam scanner with a pinhole of 1 airy unit. Cells labeled with Cy3 or Cy3-HTL were imaged using excitation from a 561 nm pulsed diode laser (40 MHz) at 10% power (0.2 mW) while collecting fluorescence emission in a 575-625 nm window. Cells labeled with Cy5 and Cy5-HTL were imaged with a 640 nm pulsed diode laser (40 MHz) at 10% power (0.2 mW) while detecting fluorescence in a 650-700 nm window. For all measurements, line scanning mode was employed with a pixel dwell time of 10 *µ*s. For fluorescence lifetime microscopy, the output from the detectors of the confocal microscope was connected to a Time Correlated Single Photon Counting (TCSPC) module (Becker and Hickl). The FLIM images were obtained using SPC-IMAGE-NG software provided by Becker and Hickl with a collection time of 30 seconds per image. Images were segmented by pixel value and processed to measure the pixel decay curves and Phasor analysis in the SPCImage software.

#### Time-lapse imaging of HeLa cells

Imaging was performed using an Andor Revolution XD spinning disk confocal microscope equipped with a 60x (1.49 N.A.) oil immersion objective. The fluorescence was collected using an Electron multiplying charge coupled device (EMCCD) (iXon 887, Andor). Samples were imaged in two channels (bright field and 607/36 nm) for PEPCy3, while being illuminated using a 561 nm laser. HeLa cells expressing PEPCy5-B2AR-Ala were excited using a 640 nm laser and imaged in two channels (bright field and 700/75) for a total duration of 30 mins.

#### Single molecule localization microscopy of HEK293 and *E. coli* cells

Prior to single molecule imaging PEPCy3-Cy3 signal was confirmed by diffraction limited wide-field epifluorescence microscopy, performed on a Nikon Eclipse Ti2 microscope equipped with a super apochromat objective (PlanApo, 100x, 1.45 N.A., oil immersion, Nikon) and scientific cMOS camera (Prime95B, Photometrix). Cells were illuminated for 100 ms with 20% of 555/28 LED light source from SPECTRA III, VBCTGYRnIR Light Source. The fluorescence emission from the sample passed through a dichroic (FF01-432/515/595/681/809-25nm, Semrock, Rochester, NY) and band pass emission filters for Cy3 (Cat. No. ET607/36, Brightline, Semrock). Single molecule localization microscopy was performed on a Nikon Eclipse Ti2 microscope using a 100x (1.45 N.A.) oil immersion objective. This setup is capable of objective based total internal reflection (TIR) microscopy. To stimulate the samples, a laser with an excitation wavelength of 561 nm or 638 nm with an intensity of approximately 500 W/cm^2^ was employed. TIR was achieved by adjusting the angle of the laser beam on the back focal plane. The emitted fluorescence from the samples underwent filtering using a multiband dichroic mirror and specific Cy3 or Cy5 emission filters (607/36 or 700/75). Finally, the fluorescence images were captured using an EMCCD (iXon 887, Andor). A total of 1000 frames were recorded for a 512-by-512-pixel area, with each frame lasting 50 ms under continuous acquisition. The single molecule data were analyzed using the Thunderstorm[27] plugin in Fiji[28]. The two channel localizations were registered using calibration images of multicolor 0.1 *µ*m fluorescent fiduciaries (Tetraspeck, ThermoFisher scientific) via affine transformation followed by single molecule tracking using the Trackmate[29] plugin in Fiji. The median speed and jump lengths were plotted using custom Python scripts.

## Back Matter

### Supplementary information

Supplementary figures (1-2) and supplementary videos (1-6) are included.

## Acknowledgments.

This work is dedicated to Prof. Marcel Bruchez and Prof. Alan Waggoner (Carnegie Mellon University). We thank Prof. Bruce Armitage (Carnegie Mellon University), Dr. Petar Petrov (UC Berkeley) and Prof. W.E. Moerner (Stanford University) for invaluable inputs. We thank Prof. David Gresham and Ina Suresh (New York University) for help with constructing the yeast strains. We thank Prof. Lucien Weiss (Polytechnique Montŕeal) for helpful suggestions on the manuscript.

- Availability of data and materials: The data reported in this work will be made available via NYU UltraViolet repository. Accession codes will be available before publication. Plasmids will be made available via Addgene.
- Code used in this work is available through the Saurabh lab Github repository https://github.com/saurabhLabNYU/PEPCy_2024

## Supplementary Figures

**Fig. S1:**
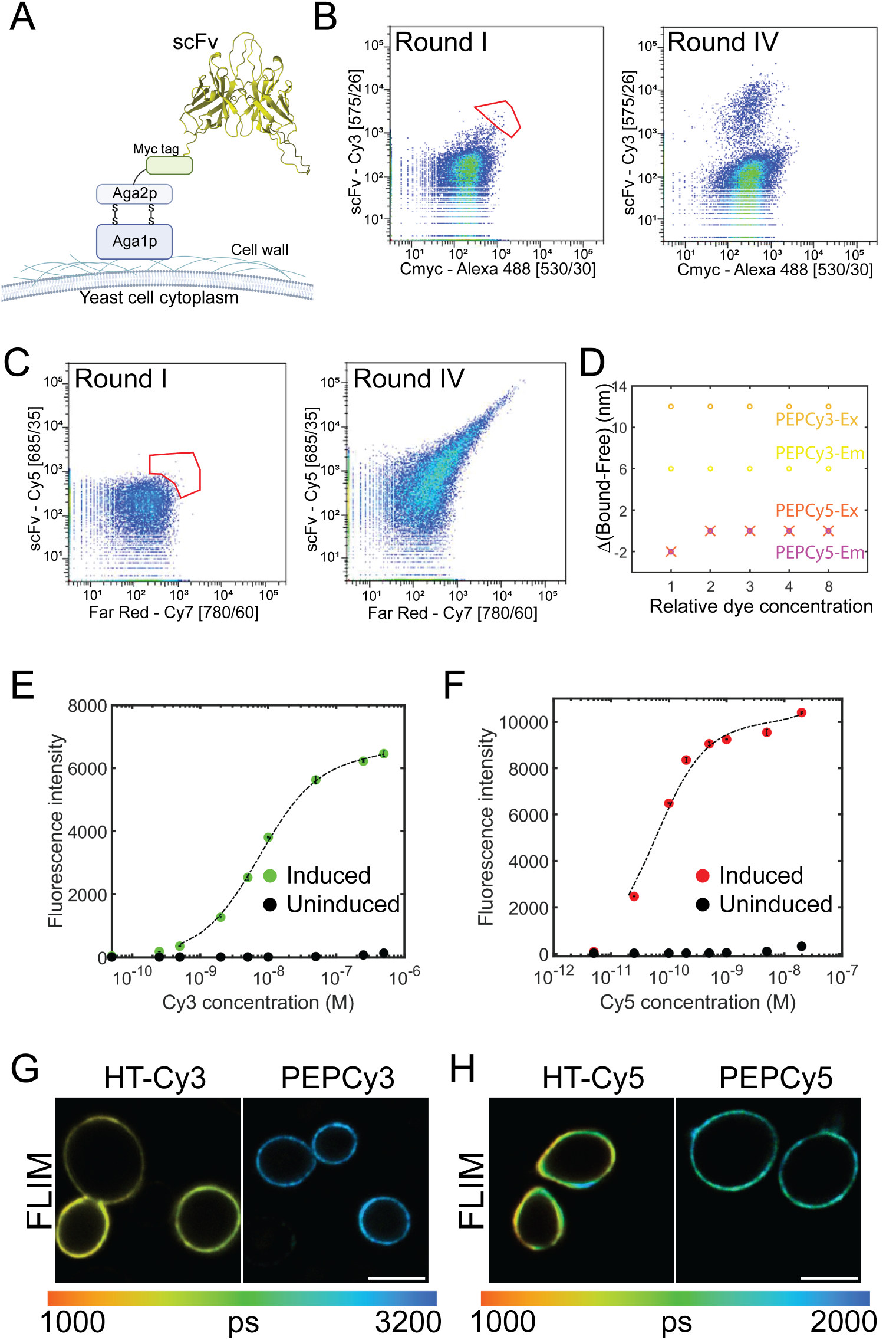
Directed evolution, spectral and binding properties of PEPCy tags. **A**. Schematic of the scFv membrane anchoring in the yeast cell surface display **B**. Distribution of cell fluorescence signal collected in the Cy3 channel and Alexa488 channel via flow cytometry. The red outline in round I indicates the gating for flow-sorting. In round IV a the population is enriched in Cy3 binders, evident from the shift in the distribution on Cy3 axis. **C**. Distribution of cell fluorescence signal collected in the Cy5 channel and Cy7 channel via flow cytometry. The red outline in round I indicates the gating for flow-sorting. In round IV a the population is enriched in Cy5 binders, evident from the shift in the distribution on Cy5 and Cy7 axes. **D**. The difference between excitation maximum or emission maximum (in nm) for free cyanine dyes and their complexes with PEPCy tags. For PEPCy3:Cy3 complex, dye concentrations of 100 nM to 800 nM were used with ten-fold excess of PEPCy3. For the PEPCy5-Cy5 complex, Cy5 was taken up from 50 pM to 400 nM with ten-fold protein excess. **E**. Binding isotherm for PEPCy3 expressed on yeast cell surface with increasing concentrations of Cy3. To account for non-specific binding, control yeast cells not expressing PEPCy were used. Each point is an average from three replicated with standard error of the mean shown over. The dashed line shows the fit of the background subtracted signal with a model for single site binding. **F**. Binding isotherm for PEPCy5 expressed on yeast cell surface with increasing concentrations of Cy5 is shown. Uninduced cells were used to estimate non-specific binding and the data were fit using a single binding site model. Each point is an average from three replicated with standard error of the mean shown over. **G**. FLIM micrographs of yeast cells expressing HT or PEPCy3 on their surface, labeled with their cognate Cy3 dyes. The images are color coded based on lifetime in picoseconds shown in the colorbar below. **H**. FLIM micrographs of yeast cells expressing HT or PEPCy5 on their surface, labeled with their cognate Cy5 dyes. The images are color coded based on lifetime in picoseconds shown in the colorbar below. Scale bars in both E and F are 5 *µ*m. A was created with BioRender.com

**Fig. S2:**
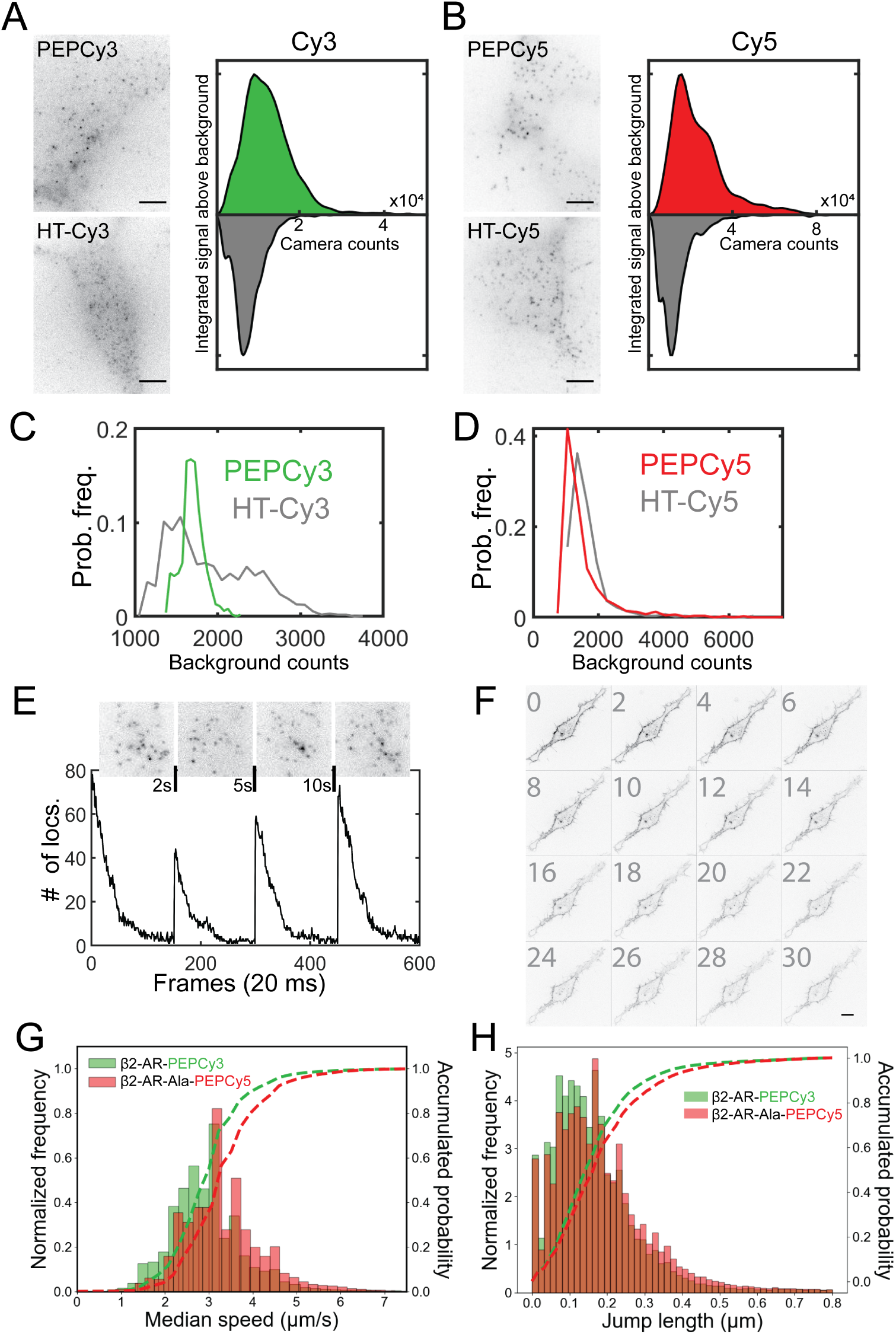
Application of PEPCy tags for single molecule and time-lapse microscopy of cell surface proteins. **A.** Micrographs of HEK cells expressing PEPCy3 or HT on their surfaces labeled with Cy3, and **B.** cells expressing PEPCy5 or HT on their surfaces labeled using Cy5. The colormap is inverted to clearly show single molecules against the white background. Normalized probability distributions of integrated single molecule signal above the background are shown adjacent to each image. (N = 1200-2000 single molecules per sample for A and B). Scale bar is 5 *µ*m. **C**. Probability distributions of the background counts from single PEPCy3 or HT-Cy3 molecules expressed on HEK cell surfaces and imaged via TIRF microscopy (N = 998 for PEP and 1020 for HT) **D**. Probability distributions of the background counts from single PEPCy5 or HT-Cy5 molecules expressed on HEK cell surfaces and imaged via TIRF microscopy (N = 1133 for PEP and 784 for HT). **E**. Wash-free extended tracking of single PEPCy5-B2AR-Ala molecules on HEK cell surface. The first image of each acquisition is shown. The black vertical lines represent the instance and duration of the illumination turned off. The graph below shows the number of localizations per frame as a function of time in this experiment. **F**. Time-lapse confocal microscopy of HeLa cells expressing B2AR-Ala-PEPCy5 labeled using Cy5. Numbers on the left of each image denote the time in minutes. Images were acquired every 30 s for 30 mins. **G**. Normalized frequency distributions (bars) and cumulative probability (dashed lines) of median speed per trajectory for PEPCy3-B2AR and PEPCy5-B2AR-Ala molecules (trajectory N = 42906 for PEPCy3 and N = 25253 for PEPCy5) **H**. Normalized frequency distributions (bars) and cumulative probability (dashed lines) of jumps in a 50 ms duration for PEPCy3-B2AR and PEPCy5-B2AR-Ala molecules.

## Supplementary video captions

**S1** - Single-molecule imaging of surface-expressed PEPCy or HT on HEK293 cells labeled via cognate Cy3 dyes. Images were acquired using a TIRF microscope at a frame rate of 20 Hz and with constant laser exposure. The movies are played in real time.

**S2** - Single-molecule imaging of surface-expressed PEPCy or HT labeled via cognate Cy5 dyes. Images were acquired using a TIRF microscope at a frame rate of 20 Hz and with constant laser exposure. The movies are played in real time.

**S3** - Wash-free, long term single-molecule imaging of PEPCy5-tagged surface receptors on HEK293 cells incubated with 10 pM Cy5 and imaged via TIRF microscopy. Each phase is 150 frames of 50 ms exposure each with the excitation laser turned off for 2 s between phase 1 and 2, for 5 s between phase 2 and 3, and for 10 s between phases phase 3 and 4.

**S4** - Time-lapse imaging of PEPCy3-B2AR expressing HeLa cells incubated with the agonist Isoproterenol followed by wash out at time t = 0. Each frame was captured with an exposure of 100 ms with a 30 s interval between frames and a total acquisition time of approximately 40 mins.

**S5** - Time-lapse imaging of PEPCy5-B2AR-Ala expressing HeLa cells in the absence of any agonist. Each frame was captured with an exposure of 100 ms with a 30 s interval between frames and a total acquisition time of approximately 30 mins.

**S6** - Simultaneous tracking of PEPCy3-B2AR and PEPCy5-B2AR-Ala receptors on HEK293 cell surface via TIRF microscopy. PEPCy3 tracks are shown in green and PEPCy5 tracks are shown in red.

## Notes

### Competing Interest Statement

The authors have declared no competing interest.

## References

[1] Grimm, J. B. & Lavis, L. D. Caveat fluorophore: an insiders’ guide to small-molecule fluorescent labels. Nat Methods 19, 149–158 (2022).

[2] Demchenko, A. P. Photobleaching of organic fluorophores: quantitative characterization, mechanisms, protection. Methods Appl. Fluoresc. 8, 022001 (2020).

[3] Grimm, J. B. et al. A general method to improve fluorophores for live-cell and single-molecule microscopy. Nat Methods 12, 244–250 (2015).

[4] Grimm, J. B. et al. A general method to fine-tune fluorophores for live-cell and in vivo imaging. Nat Methods 14, 987–994 (2017).

[5] Grimm, J. B. et al. Optimized Red-Absorbing Dyes for Imaging and Sensing. J. Am. Chem. Soc. 145, 23000–23013 (2023).

[6] Mujumdar, R. B., Ernst, L. A., Mujumdar, S. R., Lewis, C. J. & Waggoner, A. S. Cyanine dye labeling reagents: Sulfoindocyanine succinimidyl esters. Bioconjugate Chem. 4, 105–111 (1993).

[7] Vogelsang, J., Cordes, T., Forthmann, C., Steinhauer, C. & Tinnefeld, P. Controlling the fluorescence of ordinary oxazine dyes for single-molecule switching and superresolution microscopy. Proceedings of the National Academy of Sciences 106, 8107–8112 (2009).

[8] Altman, R. B. et al. Cyanine fluorophore derivatives with enhanced photostability. Nat Methods 9, 68–71 (2011).

[9] Altman, R. B. et al. Enhanced photostability of cyanine fluorophores across the visible spectrum. Nat Methods 9, 428–429 (2012).

[10] Tinnefeld, P. & Cordes, T. ’Self-healing’ dyes: intramolecular stabilization of organic fluorophores. Nat Methods 9, 426–427 (2012).

[11] Blanchard, S. C. Reply to”’Self-healing’ dyes: intramolecular stabilization of organic fluorophores”. Nat Methods 9, 427–428 (2012).

[12] Gidi, Y. et al. Unifying Mechanism for Thiol-Induced Photoswitching and Photostability of Cyanine Dyes. J. Am. Chem. Soc. 142, 12681–12689 (2020).

[13] Saurabh, S. & Bruchez, M. P. *Targeting Dyes for Biology* From Biochemistry to Nanoscopy (CRC Press, 2014).

[14] Szent-Gyorgyi, C. et al. Fluorogen-activating single-chain antibodies for imaging cell surface proteins. Nat Biotechnol 26, 235–240 (2008).

[15] Shank, N. I., Pham, H. H., Waggoner, A. S. & Armitage, B. A. Twisted cyanines: A non-planar fluorogenic dye with superior photostability and its use in a protein-based fluoromodule. J. Am. Chem. Soc. 135, 242–251 (2013).

[16] Zhang, M. et al. Fluoromodule-based reporter/probes designed for in vivo fluorescence imaging. J Clin Invest 125, 3915–3927 (2015).

[17] Tebo, A. G. et al. Orthogonal fluorescent chemogenetic reporters for multicolor imaging. Nat Chem Biol 17, 30–38 (2021).

[18] Saurabh, S., Perez, A. M., Comerci, C. J., Shapiro, L. & Moerner, W. Super-resolution imaging of live bacteria cells using a genetically directed, highly photostable fluoromodule. J. Am. Chem. Soc. 138, 10398–10401 (2016).

[19] Saurabh, S., Zhang, M., Mann, V. R., Costello, A. M. & Bruchez, M. P. Kinetically Tunable Photostability of Fluorogen-Activating Peptide-Fluorogen Complexes. Chemphyschem 16, 2974–2980 (2015).

[20] Stennett, E. M. S., Ciuba, M. A., Lin, S. & Levitus, M. Demystifying PIFE: The Photophysics Behind the Protein-Induced Fluorescence Enhancement Phenomenon in Cy3. J. Phys. Chem. Lett. 6, 1819–1823 (2015).

[21] Ploetz, E. et al. A new twist on PIFE: photoisomerisation-related fluorescence enhancement. Methods Appl Fluoresc 12, 012001 (2024).

[22] Cao, T. T., Deacon, H. W., Reczek, D., Bretscher, A. & von Zastrow, M. A kinase-regulated PDZ-domain interaction controls endocytic sorting of the beta2-adrenergic receptor. Nature 401, 286–290 (1999).

[23] Loya-Lopez, S. I. et al. Intranasal CRMP2-Ubc9 inhibitor regulates Na V 1.7 to alleviate trigeminal neuropathic pain. Pain 165, 573–588 (2024).

[24] Briggs, M. S., Burns, D. D., Cooper, M. E. & Gregory, S. J. A ph sensitive fluorescent cyanine dye for biological applications. Photochem. Photobiol. Sci. 1, 457–462 (2001).

[25] Chao, G. et al. Isolating and engineering human antibodies using yeast surface display. Nat Protoc 1, 755–768 (2006).

[26] Bacon, K. et al. Isolation of Chemically Cyclized Peptide Binders Using Yeast Surface Display. ACS Comb. Sci. 22, 519–532 (2020).

[27] Ovesný, M., Ǩŕı̌zek, P., Borkovec, J., Švindrych, Z. & Hagen, G. M. Thunder-STORM: a comprehensive ImageJ plug-in for PALM and STORM data analysis and super-resolution imaging. Bioinformatics 30, 2389–2390 (2014).

[28] Schindelin, J., et al. Fiji: an open-source platform for biological-image analysis. Nat Methods 9, 676–682 (2012).

[29] Ershov, D. et al. TrackMate 7: integrating state-of-the-art segmentation algorithms into tracking pipelines. Nat Methods 19, 829–832 (2022).

